# Methacrylic acid-based biomaterials promote peripheral innervation in the subcutaneous space of mice

**DOI:** 10.1101/2022.04.03.486876

**Authors:** Alaura M. Androschuk, Theresa H. Tam, Redouan Mahou, Cheun Lo, Michael W. Salter, Michael V. Sefton

## Abstract

Peripheral nerve innervation is essential for regulating tissue repair and regeneration. MAA-based biomaterials have been previously shown to promote angiogenesis. Here we show a new role for MAA-based biomaterials in promoting terminal axon nerve growth. Our results demonstrate that MAA-based biomaterials promote peripheral nerve growth in an *Igf-1* and *Shh* dependent manner. The resulting nerves increased the sensitivity of treated mice paws to nociception. iDISCO clearing showed that MAA increased the presence of peripheral nerve structures in whole explants. MAA was also able to increase the expression of key neuronal markers and growth factors in a peripheral neuropathy model, the diabetic db/db mouse, suggesting that MAA-based biomaterials may be relevant to treatment of peripheral neuropathy. Moreover, in a peripheral neuropathy model, MAA was able to up-regulate the expression of growth factors for an extended duration suggesting MAA may prevent degeneration through an effect on factors that promote survival. As all tissues are innervated, MAA-based biomaterials could have broad applications in the promoting regeneration and preventing degeneration of peripheral nerves.

**HIGHLIGHTS:**

- Methacrylic acid-based biomaterials promote axon growth in-situ without exogenous growth factors or cells
- Methacrylic acid-based biomaterial induced terminal axon growth displays nociception, an indicator of functional outgrowth
- Methacrylic acid-based biomaterials terminal axon growth is Igf-1 and Shh driven

## INTRODUCTION

Innervation is essential for the development, modulation, and regeneration of tissues and organs[1]. More importantly, every tissue is innervated. Peripheral nerves mediate the regenerative capacity of all tissues, and their absence completely hinders regeneration[2-5]. To promote nerve growth, therapeutic approaches have focused on increasing nerve growth through the use of exogenous growth factors (e.g. neurotrophin growth factors) and cells (e.g. Schwann cells/stem cells)[6]. However, these approaches have been unsuccessful due to inherent caveats, such as short half-lives and immunological responses[6,7]. Rather than promoting regeneration with exogenous factors and cells, we hypothesize that their production and recruitment can be promoted in situ.

Methacrylic acid (MAA)-based materials are vascular regenerative and promote angiogenesis without the need for exogenous factors (e.g. cells or growth factors)[8,9]. MAA-based biomaterials are available in multiple forms including beads[10,11], smooth coatings[12-14], and hydrogels[15], all of which display similar bioactive properties. In addition to promoting vessel formation, MAA-based biomaterials promote wound healing[9,16-18], mitigate pro-inflammatory cytokine expression[8,12,14], and polarize macrophages to a ‘reparative’ phenotype[11]. The effects on vascularization have lead to a focus on subcutaneous islet delivery[15], muscle regeneration[19] and soft tissue repair[14]. The increased expression of insulin-like growth factor (IGF-1)[11] nerve growth factor (NGF)[13], sonic hedgehog (Shh)[10,17], and microtubule-associated protein 1B (MAP1B)[13], all of which are important for peripheral nerve growth and regeneration, suggest that MAA-based biomaterials may promote nerve growth.

We report here that MAA-based biomaterials up-regulate the expression of key peripheral nerve markers (*βIII-tubulin* and *Uchl1*) and key nerve growth factors (*Igf-1* and *Shh*) at 21-35 days in a *Igf-1* and *Shh* dependent manner. Whole tissue imaging (iDISCO) confirmed the increase in nerve presence (βIII-tubulin and PGP9.5 positive regions) following treatment with MAA. *Ex vivo* experiments on dorsal root ganglion (DRG) organotypic explants demonstrated that treatment with MAA promotes axon outgrowth. We also demonstrated that MAA-based biomaterials are relevant in the context of peripheral neuropathy. MAA was able to increase the expression of key neuronal markers and growth factors in a peripheral neuropathy model (db/db) relative to controls. This sustained increased expression suggests a possible role for MAA in the context of injury wherein MAA prevents axon loss by up-regulating key factors that promote survival.

## MATERIALS AND METHODS

### Animals

All animal experiments and surgeries were conducted with the approval of the University of Toronto Animal Care Committee. Animals were housed under sterile conditions in the University of Toronto’s Division of Comparative Medicine. Mice had access to food and water ad libitum. CD1 mice (4-6 weeks), purchased from Charles River Laboratory. BKS.Cg-Dock7^m^ +/+ Lepr^db^/J (JAX Stock no. 000642) mice (8-9 weeks), both a/a + Lepr^db^/+ Lepr^db^ (db/db) and a/a Dock7^m^ +/+ Lepr^db^ (db/+) were purchased from Jackson Laboratory.

### Preparation of MAA/MM co-IDA Coated Disks

MAA or methyl methacrylate (MM) was copolymerized with isodecyl acrylate (IDA) using benzoyl peroxide (1% wt) to produce 40% copolymers (mol% of MAA)[12]. MAA or MM co-polymers were dissolved in tetrahydrofuran (50 mg/ml) and cast onto 3 or 4 mm diameter silicone disks (punched from 0.01000 thick non-sterile Long-Term Implantable Silicone Sheeting, Pillar Surgical Inc., La Jolla, CA, USA). MM and uncoated silicone disks were used as controls for all studies. MM, although similar to MAA, lacks methacrylic acid and does not have any reported angiogenic effect. Prior to implantation of MM or MAA coated disks, disks were washed twice in 70% ethanol for 20 minutes and twice in PBS for 20 minutes to sterilize.

### Preparation of Methacrylic Acid (MAA) Hydrogels

Non-degradable MAA-based hydrogels are semi-interpenetrating polymer networks (SIPN) created by reacting a blend of vinyl sulfone terminated polyethylene glycol (PEG-VS) (20 kDa, JenKem Technology USA Inc) and sodium polymethacrylate (PMAA-Na) (100 kDa; Polysciences) with dithiothreitol (DTT) (Sigma) in a Michael-type reaction as previously described[15]. Three hydrogel formulations were prepared and used for all experiments. PP8020 is a SIPN composed of 80% mol ethylene glycol and 20% mol sodium methacrylate]. PP8020 has angiogenic properties and iwas the ‘treatment’ condition for all studies. PP9010 is a SIPN composed of 90% mol ethylene glycol and 10% mol sodium methacrylate. PEG is the 3D network made from PEG-VS and DTT in the absence of PMAA-Na. PP9010 and PEG were used as controls as neither has any reported angiogenic properties. All components of hydrogels were endotoxin free and sterile filtered using a 0.22μm filter (0.2 μm Millex-GP) prior to injection into animals.

### Subcutaneous Hydrogel Injection and MAA/MM co-IDA Coated Disk Implantation

PP9010 (N=12), PP8020 (N=12), and PEG (N=12) hydrogels were injected into the dorsal subcutaneous space of CD1 mice (6-7 week old; Charles River). Briefly, CD1 mice were anaesthetized with 0.5% w/v isofluorane prior to surgery and Ketoprofen (5 mg/kg) was administered intraoperatively. Hair was removed from the dorsal flank of the mouse using a depilatory cream (Veet). Approximately 50µL of hydrogel was subcutaneously injected using a 0.5mL insulin syringe (BD Canada, Mississauga, ON) on the left and right dorsal sides of the mouse. The hydrogels were explanted at 14, 21, 28, and 35 days. Immediately following euthanization; hydrogels and surrounding tissue were excised and immediately fixed in buffered formalin for histological analysis or underwent RNA isolation for gene expression analysis.

MM-coated, MAA-coated, or uncoated silicone disks were implanted into bilateral subcutaneous pockets (one disk/pocket; two disks/animal) of each animal (N=6 per treatment group) created by blunt dissection with forceps with the same material placed in both dorsal pockets. Briefly, mice were anesthetized using 2-5% isofluorane and ketoprofen (5mg/kg) was delivered intraoperatively. On the middle of the dorsal area, a longitudinal incision 1cm in length was created using a scalpel blade. Subcutaneous pockets were created by blunt dissection with forceps. MAA-coated, MM-coated, or uncoated silicone sterile disks were inserted into the subcutaneous pockets. The incision was closed with 7.5 × 1.75 mm staples and left undressed. Staples were removed 7 days after the procedure. Immediately following euthanization, disks and surrounding tissue were excised and immediately fixed in buffered formalin for histological analysis or underwent RNA isolation for gene expression analysis.

### Inhibitor Studies

Some mice (N=3) were administered the IGF-1 inhibitor Tyrphostin AG1024 (Sigma) daily beginning on the day of hydrogel implantation for 21 days. AG1024 (30µg) was delivered in DMSO (3µL) (Sigma) and PBS (200µL) via an IP injection. Vehicle group received DMSO in PBS via an IP injection daily for 21 days. Other mice (N = 3) were administered 10mg/kg/day of Sonidegib (0.25mg; Selleck Chemicals) in 2% DMSO (Sigma) and corn oil (Sigma) for 21 days for the treatment group or 2% DMSO in corn oil for vehicle group via an IP injection. The first dose was administered on the day of hydrogel implantation and continued daily for 21 days. Hydrogels and surrounding tissues were excised and RNA was isolated.\ Samples were then assayed for gene expression via RT-qPCR.

### Subcutaneous Denervation Model

5-week old CD1 male mice underwent subcutaneous denervation surgery in which the subcutaneous nerves were removed exclusively from the right side of the animal located at anatomical sites T3-T12 using a previously established protocol[20] creating a peripheral nerve injury that is not endogenously repaired. A contralateral sham-operated control was created on the left side of the animal in which all nerves were left intact. To evaluate the effect of denervation, a hypodermic needle was used to provide stimulation to both the denervated side of the mouse and the contralateral sham. Response to physical stimuli was recorded. Some tissues were also excised and processed for histology; staining for PGP9.5 in the denervated area to confirm that the area was stably denervated at 10 days following surgery. Only mice in which the tissue had been successfully and stably denervated (based on hypodermic stimulation) underwent subcutaneous hydrogel injection to assess the effect of the MAA-based hydrogel on peripheral nerve regeneration (N=5). Mice were sacrificed and hydrogels were explanted at 21 days after implantation.

### Intraplantar Hydrogel Injection and Mechanosensation Testing

Approximately 8-week old CD-1 male mice received unilateral injections of MAA (N=6) or PEG (N=7) as control (20 μl) into the mid-plantar surface of the left hind paw. The up-down method of Dixon[21] was used to determine the 50% paw withdrawal threshold force. Mice were placed in Plexiglas cubicles on a perforated metal platform, where they were habituated for at least 1 h before testing. Calibrated von Frey monofilaments were applied to the mid-plantar surface of the hind paw and the presence or absence of withdrawal responses were recorded. If mice showed no paw withdrawal after the highest filament applied (∼1.4g), the withdrawal threshold was reported as 2g. Two consecutive measures were taken at each time point at least 15 minutes apart, and averaged.

### RNA Isolation and Gene Expression Analysis

Tissue explants (hydrogel with surrounding tissue) were placed in gentleMACS M tubes (Miltenyi Biotec Inc., Auburn, CA, US) containing lysis buffer on ice for 30 minutes. Tissue samples were homogenized using gentleMACS M tubes with the gentleMACS Dissociator (Miltenyi Biotec Inc., Auburn, CA, US). RNA isolation was completed using PuroSPIN™ Fibrous Tissue RNA Purification Kit (Luna Nanotech, Toronto, ON) or QIAGEN Fibrous Tissue Kit (LOCATION) according to manufacturer’s instructions with the exception of the proteinase K digestion step in which the tissue was digested for 15 minutes at 55ºC instead of the suggested 10 minutes. RNA quantity, quality, and purity of RNA was assessed using a NanoDrop ND-1000 spectrophotometer (NanoDrop Technologies Inc., Wilmington, DE, US). The 260nm/280nm ratio was above 2.0 for all samples. 1µg/µl of RNA was reverse transcribed into complementary DNA (cDNA) using SuperScript III First Strand Super Mix (Invitrogen, Burlington, ON) according to the manufacturer’s instructions and assayed for mRNA of interest using BioRad Ssofast Evagreen Reaction Mix (BioRad Laboratories, Mississauga, ON). Primers (Table 1) were either designed using Primer-BLAST or selected from Primerbank and then synthesized by ACGT (Toronto, ON). All quantification was performed with the BioRad CFX Manager using the ΔΔCt method normalized to GAPDH. All samples were run in triplicate.

**Table 1.**
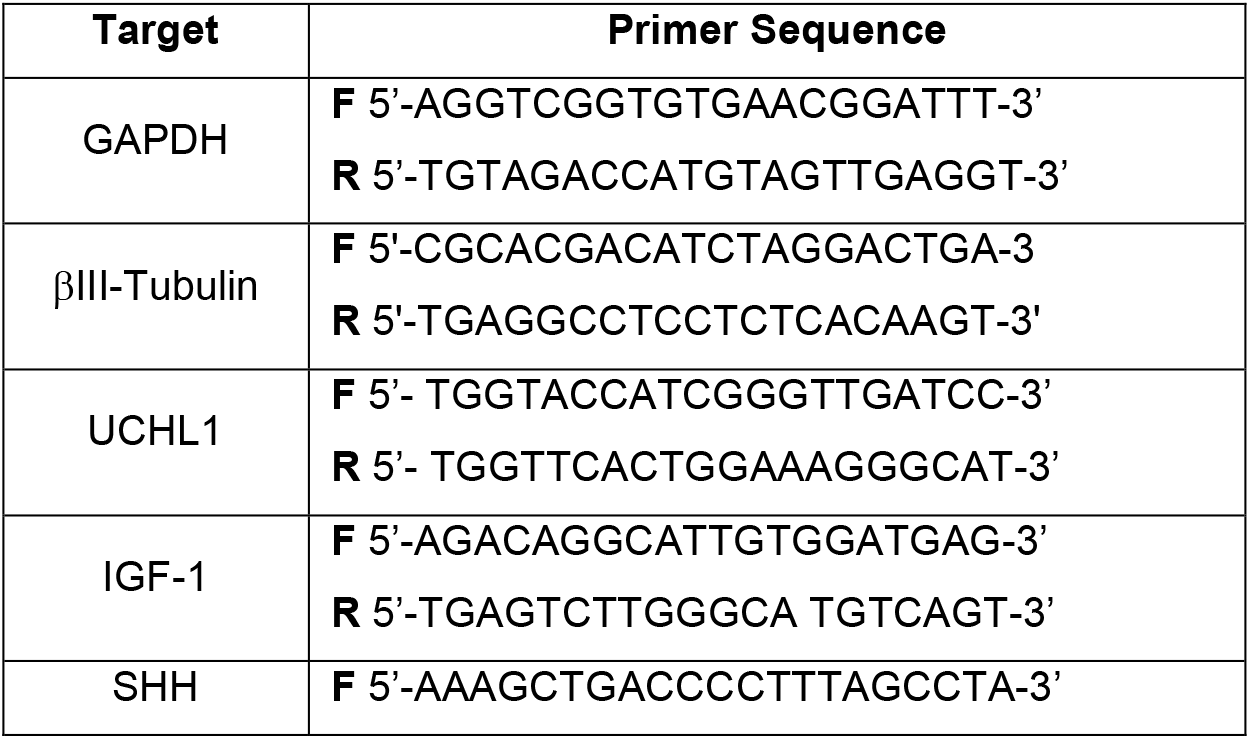
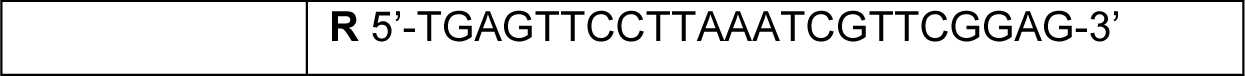
Primers used to assay the expression of peripheral nerve growth markers and growth factors

### Histology

Tissue explants were placed into a histology cassette and fixed in buffered formalin and submitted to the Pathology Research Program Laboratory (UHN, Toronto, Ontario) for immunohistochemical processing. For histological analysis, explanted tissues were embedded in histogel followed by paraffin and cut on edge in 200μm sections and stained for nerve growth (PGP9.5; Biorad). Processed slides were scanned at 20X at the Advanced Optical Microscopy Facility (MaRS, Toronto, ON) using an Aperio ScanScope XT (Leica Microsystems) and analyzed using Aperio ImageScope.

### iDISCO

iDISCO is a whole-mount, clearing and immunolabeling technique that allows for visualization of molecular markers in large, complex, intact tissues[22]. Some mice were euthanized and hydrogels and surrounding tissues were excised and post-fixed in 4% PFA in PBS at 4ºC overnight. Samples were washed thrice in PBS for 1 hour, then pre-permeabilized in 0.1% Tween-20/0.1% Triton X-100/0.1% deoxycholate/0.1% NP40/20% DMSO/PBS at 4ºC overnight. Samples were washed in PBS twice for 15 minutes followed by permeabilization in increasing concentrations of ethanol; 50%ethanol/PBS, 80%ethanol/ddH_2_O, and 100% anhydrous ethanol at 4ºC overnight. To reduce background, samples were bleached in 5%H_2_O_2_ in 20% DMSO/ethanol overnight at 4ºC followed by washing in 20% DMSO/ethanol, 80% ethanol/ddH_2_O, 50% ethanol/PBS, 100% PBS, and twice in 0.2% PBST at 4ºC for 4 hours. Samples were then incubated in 0.2% Triton X-100/20% DMSO/0.3 M glycine/PBS at 4ºC followed by blocking in 0.2% Triton X-100/6% donkey serum/10% DMSO/PBS at 37ºC.

Samples were incubated with primary antibody βIII-Tubulin (Abcam) diluted in 0.2% Tween/10µg/ml heparin/3% donkey serum/5% DMSO/PBS at 37ºC for 10 days. After incubation with primary antibody, samples were washed with 0.2% Tween-20/10µg/mL heparin/PBS 5 times for 1 hour, followed by incubation with secondary antibody (Alexa 568, Invitrogen) diluted in 0.2% Tween/10µg/ml heparin/3% donkey serum/5% DMSO/PBS at 37ºC for 10 days. Samples were then washed with 0.2% Tween-20/10µg/mL heparin/PBS 10 times for 1 hour followed by dehydration in increasing concentrations of ethanol; 50%ethanol/PBS, 80% ethanol/ddH_2_O, 100% anhydrous ethanol at 4ºC overnight. Samples were delipidized and further dehydrated in 40% 2,2,2-Trichloroethanol/40% Benzyl alcohol and then placed in 45% Diphenyl ether/30% 2,2,2-Trichloroethanol/25% Benzyl alcohol to match the refractive index for imaging. Samples were imaged using the Zeiss LSM 880 Elyra Super-Resolution Microscope at the Imaging Laboratory (University of Toronto, Toronto, ON). Images were analyzed using Imaris (version 8).

### Dorsal Root Ganglion (DRG) Isolation and MAA

Embryonic DRGs (E15) were harvested from Sprague Dawley rats and cultured with MAA or control polymers. DRGs were isolated and plated onto poly-D-lysine and laminin coated tissue culture plates for 24 hours prior to treatment with biomaterials/hydrogels to allow DRGs sufficient time to attach to the plate. Following the 24-hour attachment period, DRGs were treated with either co-polymer coated glass coverslips (40% MM co-IDA, 40% MAA co-IDA, or uncoated glass coverslips). Coated coverslips were inverted and placed on top of DRGs. During treatment, DRGs were maintained in a minimal volume of media to ensure coverslips were in contact with DRGs without damaging or preventing nutrient exchange. DRGs were treated for 24 hours and imaged using the Zeiss Live Imaging Microscope (Centre for Research and Applications in Fluidic Technologies, University of Toronto, Toronto, ON). Images were assessed using ImageJ. DRGs were maintained in fully supplemented neurobasal media (1X B27, 1X GlutaMax, 50ng/mL NGF, and 1% P/S) for all experiments.

### Statistical Analysis

Statistical analysis was performed using GraphPad Prism (Version 6). Comparison among treatment groups were performed using two-way ANOVA followed by Tukey’s post hoc test for significance. Significance was set at p<0.05. Number of biological replicates (N) is given in figure captions. All data sets represent mean ± standard error of mean (S.E.M.).

## RESULTS

### MAA-based biomaterials increased the expression of genes associated with peripheral nerve growth

In order to determine if MAA-based materials promote peripheral nerve growth, both a MAA gel and MAA coated silicone disks were tested for their effects on nerve markers.

40% MAA co-IDA, 40% MM co-IDA, or uncoated silicone disks were implanted into the subcutaneous space of CD1 mice for 21 days. Immunostaining for the peripheral nerve marker PGP9.5 showed that there was an increase in PGP9.5 positive regions in 40% MAA co-IDA treated tissue compared to 40% MM co-IDA, and untreated tissue (Fig.1A). Gene expression analysis showed that there was also an increase in the expression of *Uchl1* and *βIII-tubulin*, nerve specific markers (Fig.1B).

**Figure 1.**
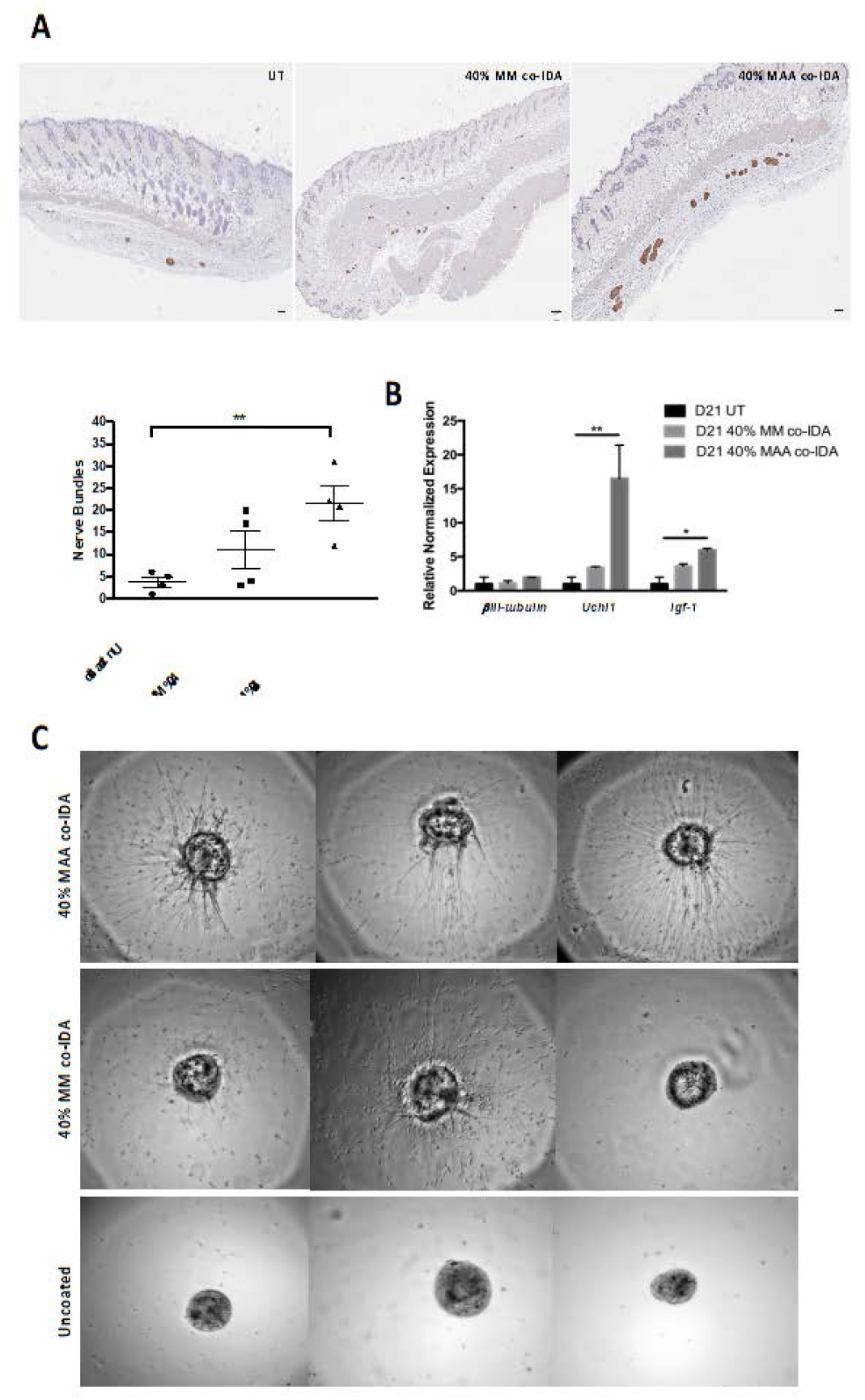
MAA-based coating promoted peripheral nerve growth in vivo and in vitro. 40% MAA-co-IDA coating of silicone disk increased the expression of the peripheral nerve markers PGP9.5, Uchl1 and βIII-tubulin in the subcutaneous space relative to untreated control (UT) or 40% MM-co-IDA coated disk. A) representative histological sections of subcutaneous tissue immunostained for the peripheral nerve marker PGP9.5, 21 days after implantation. Scale bar=100µm. More PGP9.5 positive regions were quantified in histology sections of MAA explants; counts were highly variable and this method was abandoned in favour of iDISCO images. B) qRT-PCR analysis of subcutaneous explants at D21 post-implantation. Quantification of mRNA expression was performed using ΔΔCt method relative to Gapdh and normalized to untreated samples. N=3 biological replicates for all groups; error bars indicate S.E.M. *=p<0.05;**=p<0.001 (two-way ANOVA). C) More axon outgrowth was observed in vitro with 40% MAA co-IDA treated dorsal root ganglions (DRG) following 24 hour with coated coverslips lightly sitting atop them. DRGs were isolated from E15 rat embryos and plated onto glass coverslips coated with poly-D-lysine and laminin. N=3; Each image represents a different replicate.

To mimic the effect of MAA coated disks, DRGs were isolated and cultured with 40% MAA co-IDA coated, 40% MM co-IDA coated, or uncoated coverslips for 24 hours. DRG explants contain both neurons and Schwann cells and represent a reasonable approximation to *in vivo* conditions. DRGs treated with 40% MAA co-IDA had an increase in axon length and outgrowth compared to 40% MM co-IDA and uncoated coverslips (Fig.1C), suggesting MAA promotes the outgrowth of axons consistent with the increase in expression of peripheral nerve markers seen *in vivo* with coated disks.

MAA-based hydrogels with different proportions of MAA were also injected into the subcutaneous space of mice. Gene expression analysis showed that there was an increase in the expression of *Uchl1* and *βIII-tubulin*, nerve specific markers at 21, 28, and 35 days. Expression of *Igf-1* and *Shh*, both of which are important factors for MAA-associated vascularization and perhaps also peripheral nerve growth, was also up-regulated compared to controls (Fig. 2A). There was an increase in *Igf-1* by MAA (20% MAA; PP8020) relative to controls (PP9010 (10% MAA), PEG) at day 14, but there was no up-regulation of *Uchl1* and *βIII-tubulin*. The day 14 increase in *Igf-1* is consistent with the timeline associated with MAA-induced vascularization.

**Figure 2.**
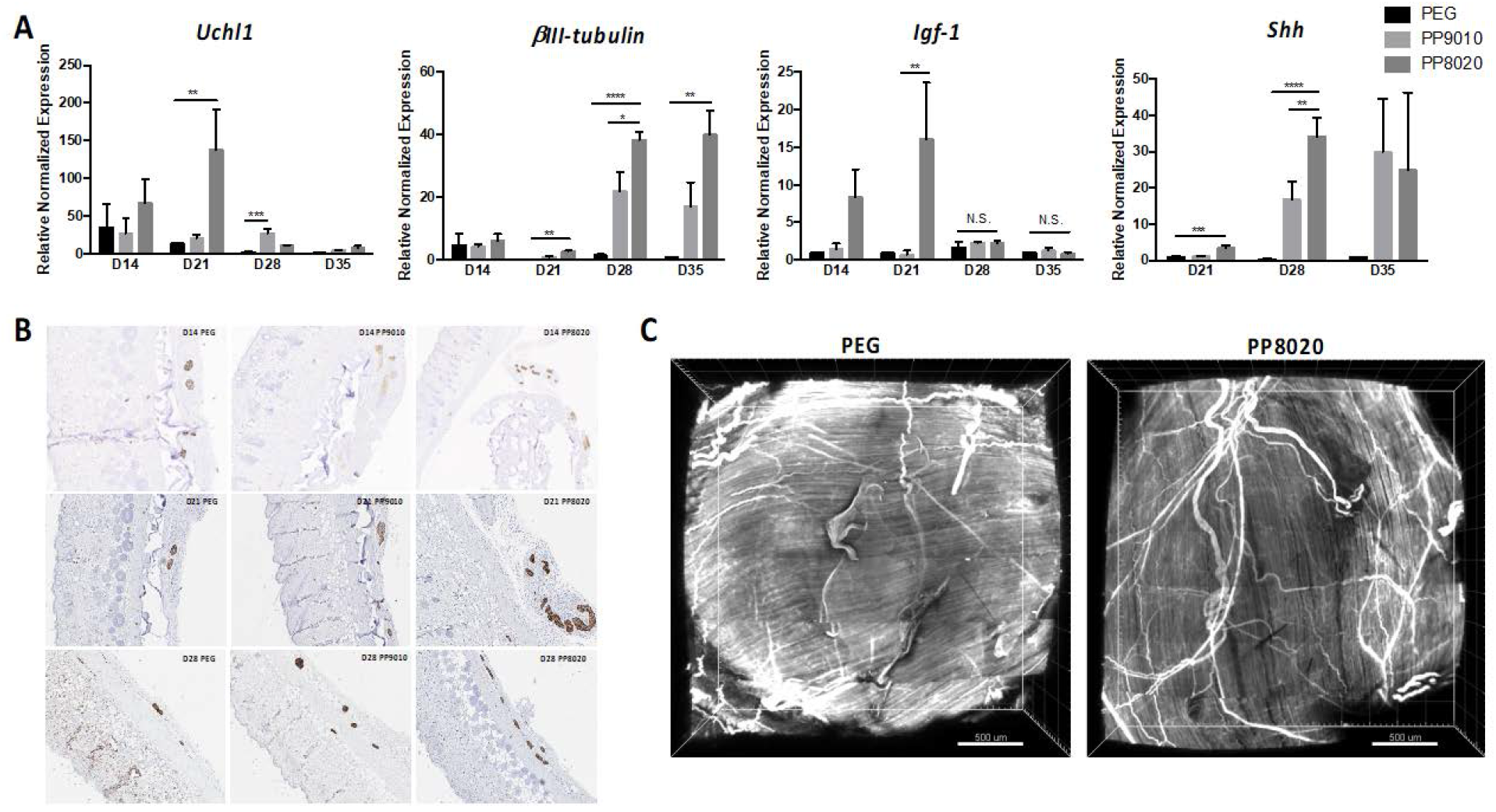
Injectable methacrylic acid semi-interpenetrating polymer network (SIPN) hydrogel promoted subcutaneous peripheral nerve growth. A) Injection of 20% MAA hydrogel (PP8020) increased the expression of Uchl1 and βIII-tubulin in the subcutaneous tissue of CD1 mice at 21, 28, and 35 days relative to that seen with a 10% MAA gel (PP9010) or a no MAA gel (PEG). PP8020 also up-regulated the expression of Igf-1 at 21 days and Shh at 21 and 35 days relative to PP9010 and PEG. mRNA quantification was performed using ΔΔCt method relative to Gapdh and normalized to PEG samples. N=3 biological replicates for all biomaterial groups and timepoints; error bars indicate S.E.M. *=p<0.05;**=p<0.001;***=p<0.0001=;****=p<0.00001 (Two-way ANOVA). B) Following subcutaneous injection of PEG, 10% MAA, or 20% MAA hydrogels, The greatest number of PGP9.5 positive regions appeared in 20% MAA subcutaneous explants at day 21 and 28 as compared to 10% MAA and PEG explants; fewer PGP9.5 positive regions appeared at day 14. PGP9.5 positive areas appear as brown regions within the tissue. Scale bar=100µm. C) A greater amount of *β*III-tubulin positive peripheral nerves were present in PP8020 subcutaneous tissues explants relative to PEG explants. Whole-mount immunolabellng of day 21 subcutaneous tissue explants from CD1 mice. Hydrogels and surrounding subcutaneous tissues were subjected to iDISCO tissue clearing and analyzed for βIII-tubulin (white). Three-dimensional volumes reconstructed from confocal micrographs of intact cleared explants. Scale bar=500µm.

Immunostaining for the peripheral nerve marker PGP9.5 shows that there was an increase in PGP9.5 positive regions in PP8020 and PP9010 compared to PEG (Fig. 2B). Gene expression analysis of pain markers (*Cox 2, Cgrp, Tac1*, and *IL-1β)* suggested that this peripheral nerve growth was not causing pain (Suppl. Fig. 1).

To validate the effect that MAA-based biomaterials have *in vivo*, the whole-mount tissue clearing technique iDISCO was employed to examine the 3D architecture of tissue and peripheral nerves following treatment. There was an increase in βIII-tubulin positive regions in PP8020-treated explants at D21 compared to PEG-treated explants, confirming that MAA gel promotes peripheral nerve growth (Fig. 2C).

### MAA-induced peripheral nerve growth exhibited nociception

To show that the MAA-induced nerve growth was functional we injected MAA/PP8020 and PEG hydrogel into the intraplantar footpad of CD1 mice. Paws/mice with intraplantar injections of MAA/PP8020 exhibited a lower von Frey withdrawal threshold than mice/paws with the PEG hydrogel injection or the contralateral paws (Fig. 3A-B), indicating increased sensitivity. This suggested that nerve growth in the MAA treated paw exhibits mechanical noncicpetion (sensitivity to mechanical stimuli).

**Figure 3.**
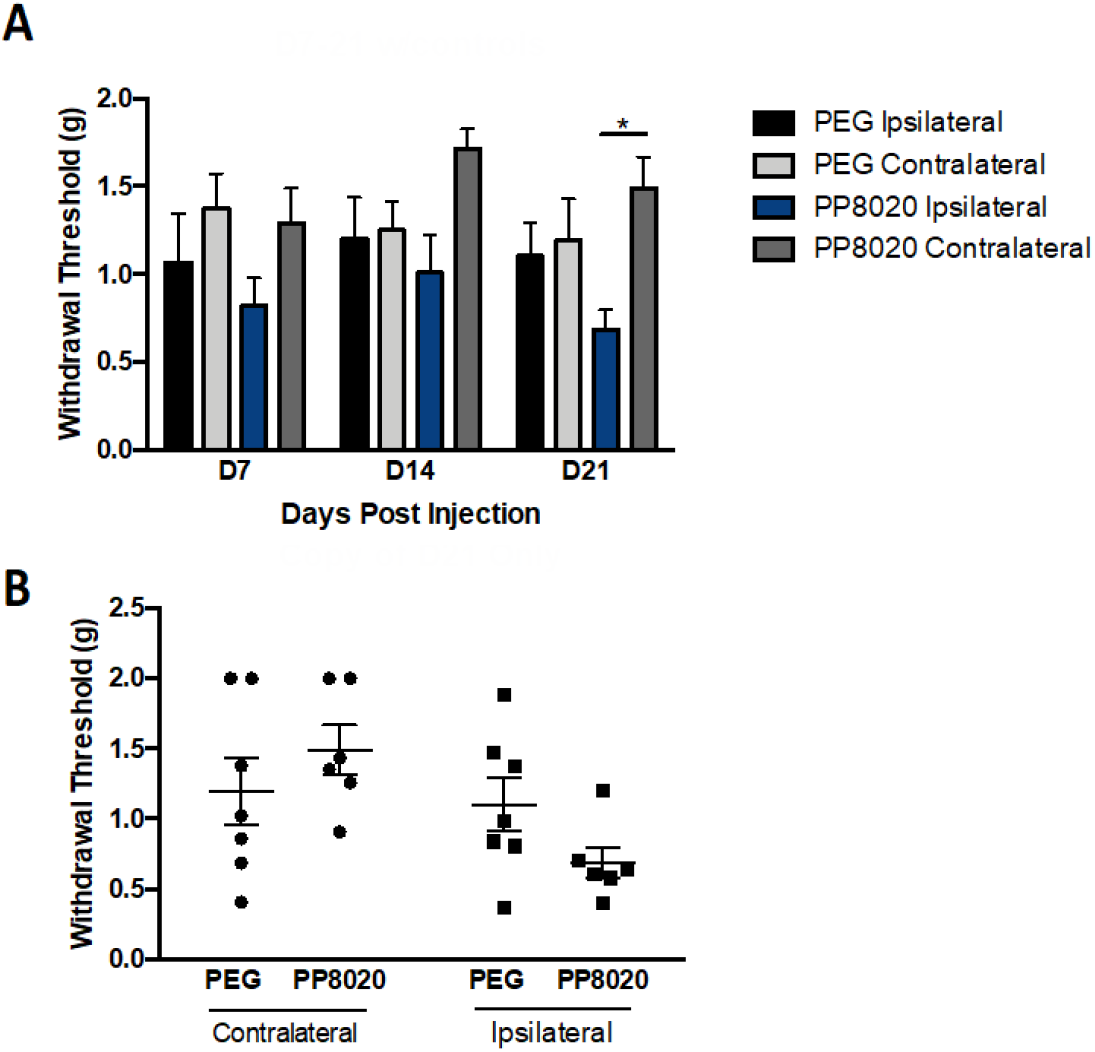
Methacrylic acid-based hydrogel induced nerve growth enhanced touch sensitivity (nociception) 20% MAA hydrogel increased paw nociception relative to PEG hydrogel. Paws with PP8020 hydrogel exhibited a lower withdrawal threshold (greater sensitivity) at 21 days (but not earlier) relative to contralateral control (no treatment paws) as well as PEG treated paws. A) Effect of implant time on average sensitivity. B) Withdrawal threshold at day 21; scatter plot showing each animal. PEG or PP8020 hydrogels were injected into the plantar surface of the paw and subjected to the Von Frey assay for nociception. Force was linearly increased and force at which paw withdrawal occurred was recorded. N=7 (PEG) and N=6 (MAA) biological replicates; error bars indicate S.E.M. *=p<0.05 (Two-way ANOVA)

Taken together these results (Figures 1-3) showed that MAA-based biomaterials do promote functional, subcutaneous peripheral nerve growth.

### MAA-based peripheral nerve growth was Igf-1 driven

Igf-1 is an important neurotrophin in nerve development, growth, regeneration, and survival[23]. Given that Igf-1 is upregulated in explanted tissues treated with MAA, we asked whether Igf-1 functions in MAA-induced peripheral nerve growth by inhibiting the Igf-1 receptor/pathway. Inhibition of Igf-1 signalling through the Igf-1R[24] resulted in the inhibition of *Igf-1 and the* decrease in expression of *βIII-tubulin, Uchl1*, and *Shh* in PP8020 treated mice relative to mice that were treated with the PEG vehicle (Fig. 4A). Immunostaining for the peripheral nerve marker PGP9.5 shows that there was a decrease in PGP9.5 positive regions in AG1024 treated PP8020 explants compared to vehicle treated PP8020 explants (Fig.4B). MAA-induced peripheral nerve growth was driven by Igf-1.

**Figure 4.**
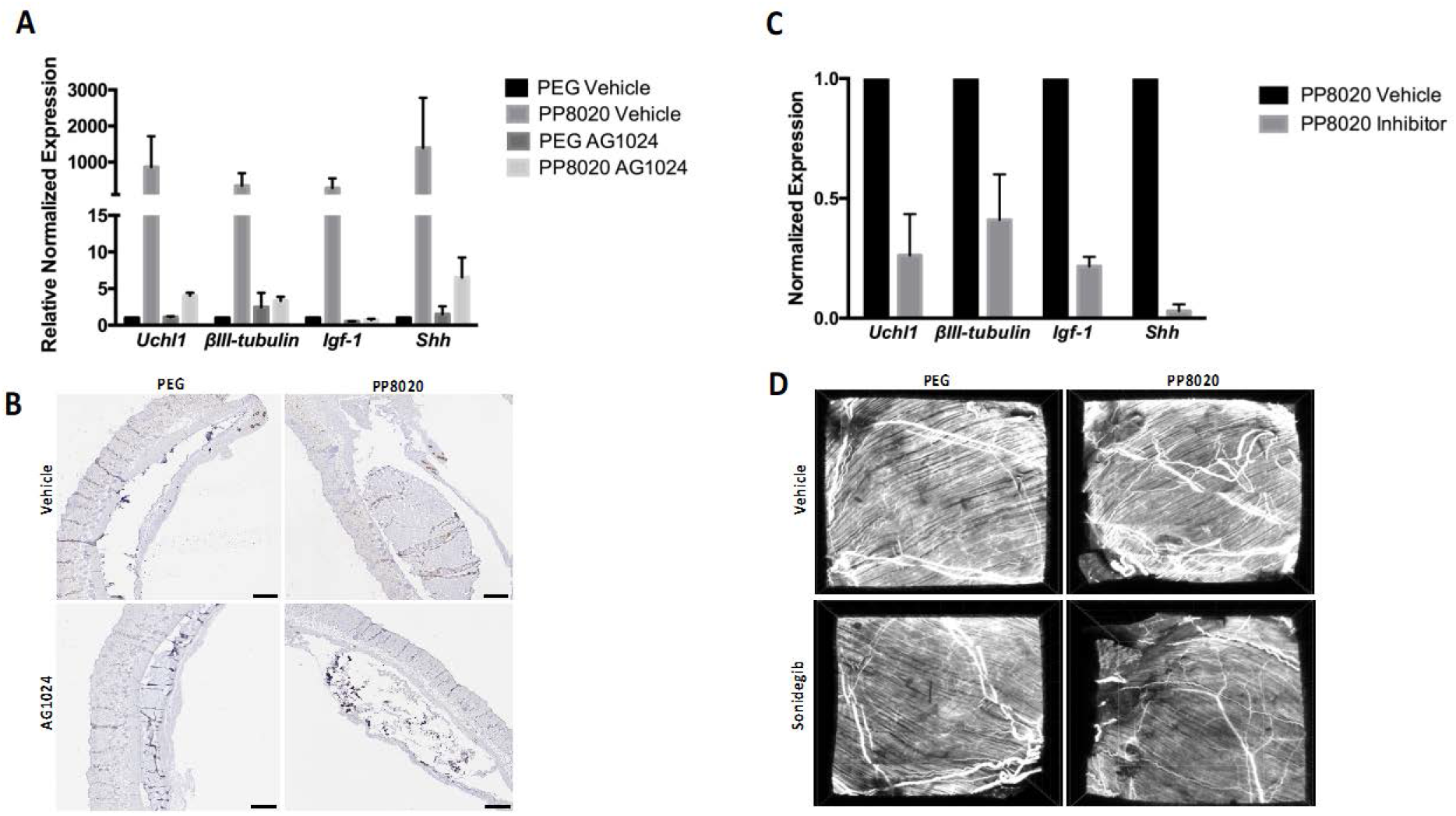
Igf-1 and Shh are required for MAA-induced peripheral nerve growth. Inhibiting Igf-1 prevented the increase in expression of the peripheral nerve markers Uchl1 and βIII-tubulin in 20% MAA subcutaneous tissue explants relative to controls. A) Inhibition of Igf-1 decreased the expression of Uchl1, βIII-tubulin, Igf-1 and Shh in PP8020 explants. qRT-PCR analysis of PEG or PP8020 subcutaneous explants following 21-day treatment with the Igf-1 inhibitor, AG1024, or vehicle.) Quantification of mRNA was as in Figure 1. N=3 biological replicates per treatment and biomaterial group; error bars indicate S.E.M. B) A decrease in PGP9.5 positive regions in 20% MAA explants was observed when Igf-1 was inhibited. Representative histological sections of subcutaneous tissue immunostained for PGP9.5 (peripheral nerve marker) of 20% MAA or PEG explants following 21-day treatment with AG1024 or vehicle. Scale bar=500µm. C) Shh inhibition resulted in the decreased expression of Uchl1, βIII-tubulin, Igf-1 and Shh in PP8020 subcutaneous explants as compared to vehicle control. qRT-PCR analysis of CD1 mice injected with PP8020 (20% MAA) hydrogel and treated with the Shh inhibitor Sonidegib for 21 days. Quantification of mRNA was as in Figure 1. N=2 or 3 biological replicates per treatment group; error bars indicate S.E.M. D) A decreased in βIII-tubulin positive peripheral nerves was observed in PP8020 subcutaneous explants when Shh was inhibited. Whole-mount immunolabellng of PEG or PP8020 (20% MAA) hydrogel explants following 21 day treatment with Sonidegib or vehicle. Subcutaneous explants were subjected to iDISCO tissue clearing and analyzed for βIII-tubulin (white). Three-dimensional volumes reconstructed from confocal micrographs; Scale bar=500µm.

### Shh was required for MAA-induced peripheral nerve growth

Shh is a morphogen required for both embryonic and adult peripheral nerve growth and regeneration[25]. Gene expression data showed that MAA hydrogels increased the expression of *Shh* at 21 and 28 days as compared to controls (Fig. 2A), which also corresponded with the increase in expression of peripheral nerve markers implying that Shh is part of the mechanism through which MAA promotes peripheral nerve growth. Therefore, we asked if Shh is required for MAA-induced peripheral nerve growth by inhibiting Shh signalling. Mice were administered Sonidegib, a smoothened antagonist[26], or vehicle for 21 days. Mice treated with MAA (PP8020) and the Shh antagonist exhibited complete inhibition in the expression of *Shh* (Fig. 4C) and the decreased expression *of βIII-tubulin, Uchl1*, and *Igf-1* compared to mice treated with MAA and the vehicle (Fig.4C).

Whole-mount tissue clearing followed immunolabeling for βIII-tubulin with volume imaging of 21-day Sonidegib and vehicle-treated PEG and PP8020 explants showed that there was no difference in the presence of βIII-tubulin positive regions between the vehicle and Sonidegib treated PEG explants (Fig.4D). On the other hand, there was a decrease in βIII-tubulin positive regions in Sonidegib-treated PP8020 explants compared to vehicle-treated PP8020 explants (Fig.4D), indicating that MAA-induced peripheral nerve growth was indeed Shh driven.

### MAA-based biomaterials promoted an increase in the expression of peripheral nerve markers in a diabetic neuropathy model

We next asked if MAA-based biomaterials could promote peripheral nerve regeneration by unilaterally denervating mice wherein nerves were removed from one side of the dorsal skin while the contralateral side remained intact as a sham internal control[20]. In this model, surgical removal of the nerve trunks resulted in the complete degeneration of the subcutaneous/cutaneous nerves leading to the loss of sensory function and indicating that skin/subcutaneous space was successfully denervated. Denervation was confirmed by sensory functional testing and immunostaining for PGP9.5, which confirmed the absence of axons in denervated tissue.

10 days following denervation PP8020, PP9010, or PEG hydrogels were injected into the denervated subcutaneous space (Fig. 5A). Analysis 21 days later showed that there was no increase in the expression of *Uchl1* and *βIII-tubulin* in denervated mice in MAA treated compared to controls (Fig. 5B). There was also no change in expression of *Igf-1* in the denervated model and no difference in response to sensory stimuli in denervated mice treated with MAA compared to controls. Immunostaining showed that there was no difference in PGP9.5 positive staining among any of the treatment groups, and a lower number of PGP9.5 positive regions compared to the contralateral innervated sham (Fig. 5C). The absence of nerve growth in the denervated model suggested that MAA may not enable de novo nerve regeneration but may promote outgrowth or the branching of existing axons in direct contact with or in close proximity to existing nerves.

**Figure 5.**
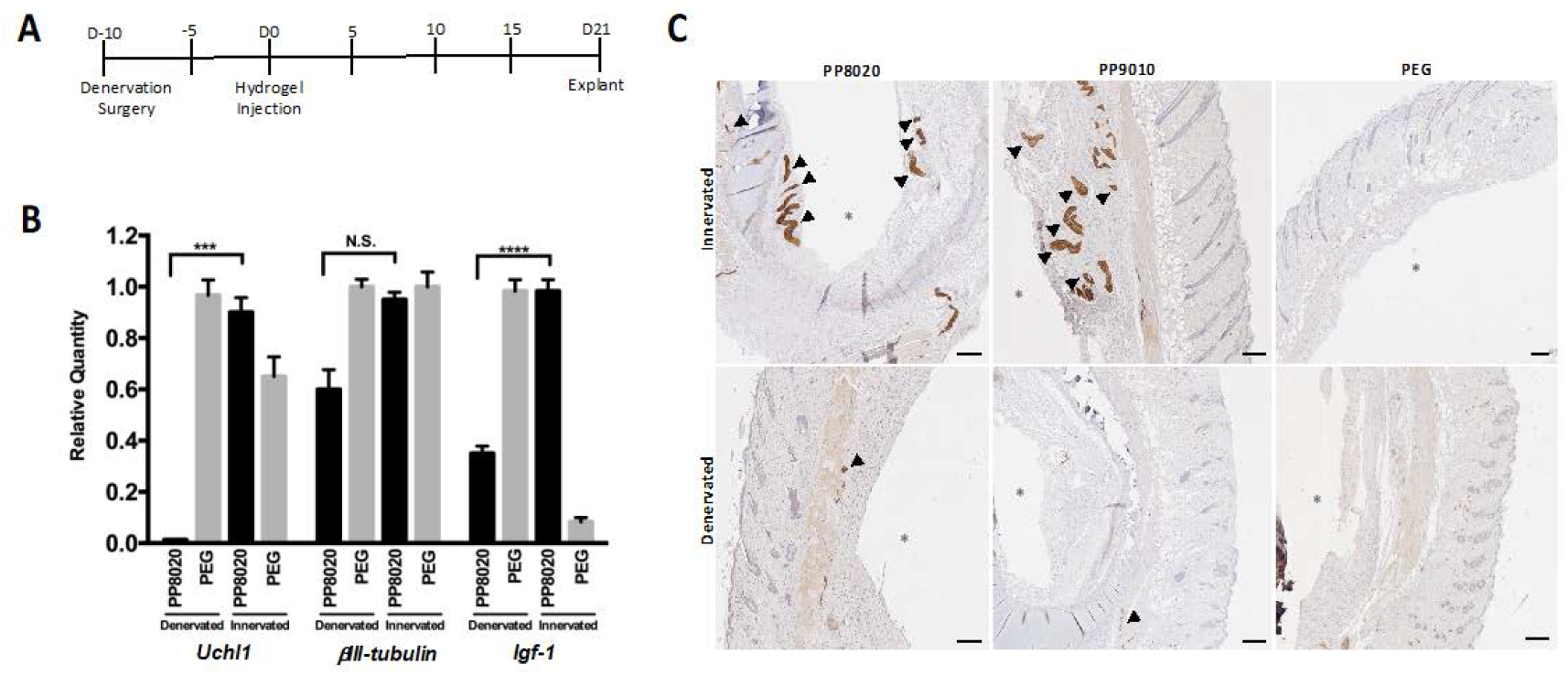
Methacrylic acid-based hydrogels did not promote de novo peripheral nerve growth in denervated subcutaneous tissue. 20% MAA did not increase the expression of peripheral nerve markers. A) Experimental timeline. Subcutaneous tissue was surgically denervated 10 days prior to hydrogel injection. Hydrogels were explanted 21 days after hydrogel injection. B) PP8020 did increased the expression of Uchl1, βIII-tubulin, or Igf-1 in the innervated model at 21 days but not when mouse was denervated. qRT-PCR analysis in innervated and denervated models after subcutaneous injection of PEG or PP8020 (20% MAA). Quantification of mRNA was as in Figure 1. N=3 biological replicates per biomaterial group per treatment group; error bars indicate S.E.M. C) No increase in PGP9.5 positive staining was observed at day 21 in denervated MAA hydrogel subcutaneous explants as compared to PEG, unlike what was seen in innervated animals (arrows denote PGP9.5 positive regions). Scale bar=200µm.

Given these findings we asked if MAA-based biomaterials could function in promoting the repair of damaged axons in a peripheral nerve injury model. Diabetic peripheral neuropathy is characterized by a decrease in systemic Igf-1 levels as well as local Igf-1 and Igf-1R expression in peripheral nerves[27,28]. Lack of Igf-1 results in axon withdrawal from terminal innervation[29,30]. Mice homozygous for the spontaneous mutation of the leptin receptor (Lepr^db^;db/db), display diabetic peripheral neuropathy[31]. To determine if MAA-based biomaterials could promote repair of damaged peripheral nerves, PP8020, PP9010, or PEG hydrogels were injected into the subcutaneous space of db/db mice at 8 weeks, at which time peripheral nerve damage has occurred[32]. We characterized the effect of the MAA hydrogels on nerve growth over time in this model. At 21 and 28 days post hydrogel injection there were slight but not significant increases in the expression of nerve markers in MAA-treated tissues relative to the control (Fig. 6A); this contrasted with the effects seen in CD1 mice (Fig 2). Only at 35 days, was there an increase in the expression of peripheral nerve markers and growth factors in MAA treated samples relative to a control (Fig. 6A). Immunostaining showed that there was an increase in PGP9.5 positive staining in PP8020 treated db/db mice compared to PEG treated controls (Fig. 6B) at day 35. This increase suggests that MAA may promote regrowth of damaged nerves in the db/db diabetic model or may prevent further axon die-back by up-regulating the expression of factors needed for neuron survival.

**Figure 6.**
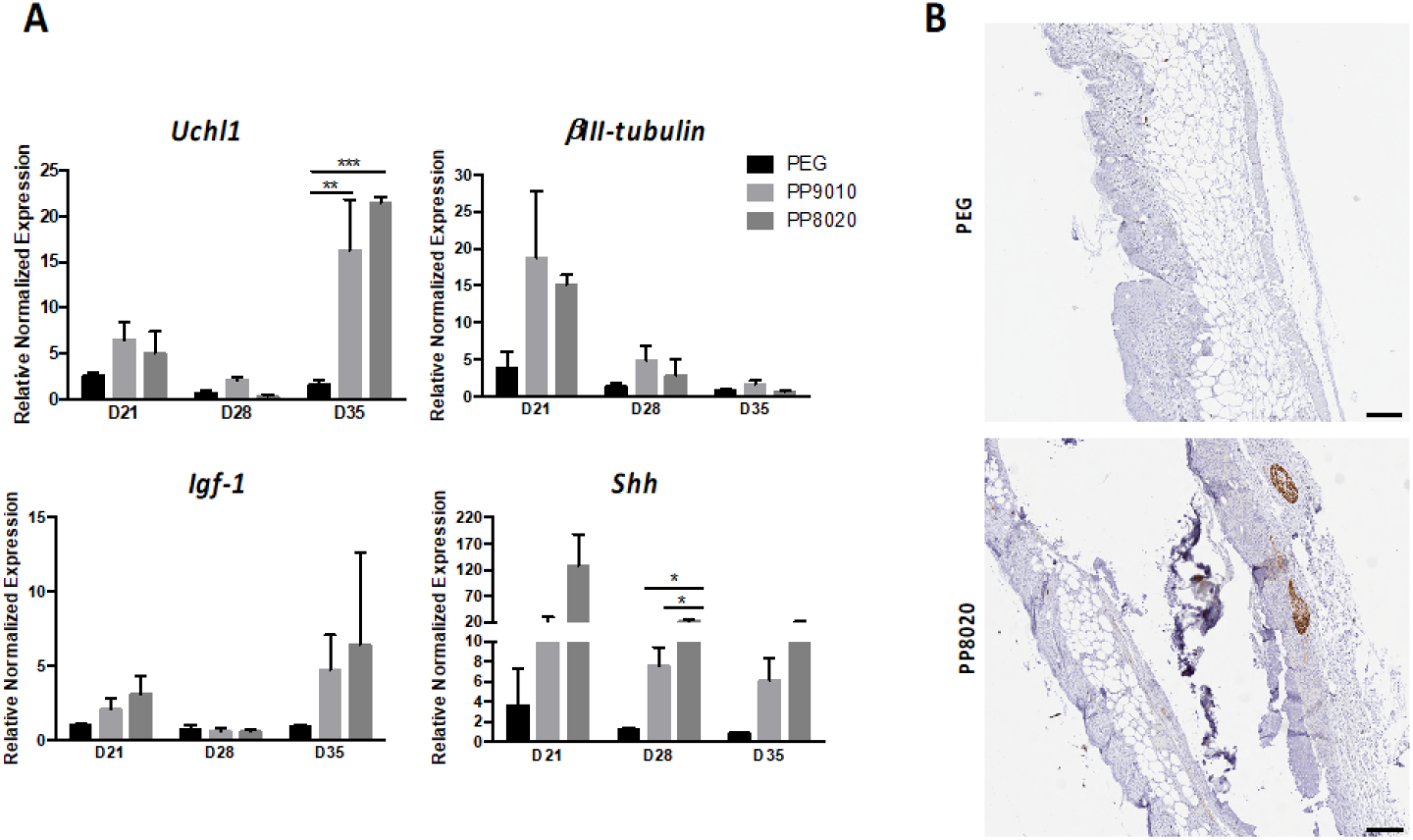
Methacrylic acid-based hydrogels promote peripheral nerve growth in db/db mice. PP8020 increased the expression of the peripheral nerve marker *Uchl1* and growth factor *Shh* in a diabetic peripheral neuropathy model relative to PEG. A) Injection of 20% MAA hydrogel increased the expression of *Shh* in the subcutaneous explants of db/db mice at 21, 28, and 35 days relative to PEG; *Uchl1* was increased at 35 days but not earlier Quantification of mRNA was performed as in Figure 1. N=3; error bars indicate S.E.M. *=p<0.05;**=p<0.001;***=p<0.0001=;****=p<0.00001 (Two-way ANOVA). B) 20% MAA increased the number of PGP9.5 positive regions at 35 days in db/db explants. Scale bar=200µm

### MAA-induced blood vessel growth is nerve dependent

Paracrine mediators from axon terminals have been shown to promote tissue regeneration and angiogenesis[2,3]. On the basis of these findings we asked whether the effect of MAA was nerve dependent. Given that MAA promotes angiogenesis, we asked if MAA-induced vessel formation was affected by the absence of nerves. To assess this, we denervated the subcutaneous space of mice as described above and immunostained for CD31-positive blood vessels at 21 days following PP8020 or PEG injection. In the absence of innervation there was a decrease in the number of CD31-positive blood vessels in PP8020-treated explants compared to PP8020-treated sham explants (Fig. 7). CD31-positive regions were not seen in PEG-treated denervated and sham explants (Fig. 7). Thus, denervation impaired the angiogenic effects of MAA as well as the nerve regeneration effects shown in Figure 5.

**Figure 7.**
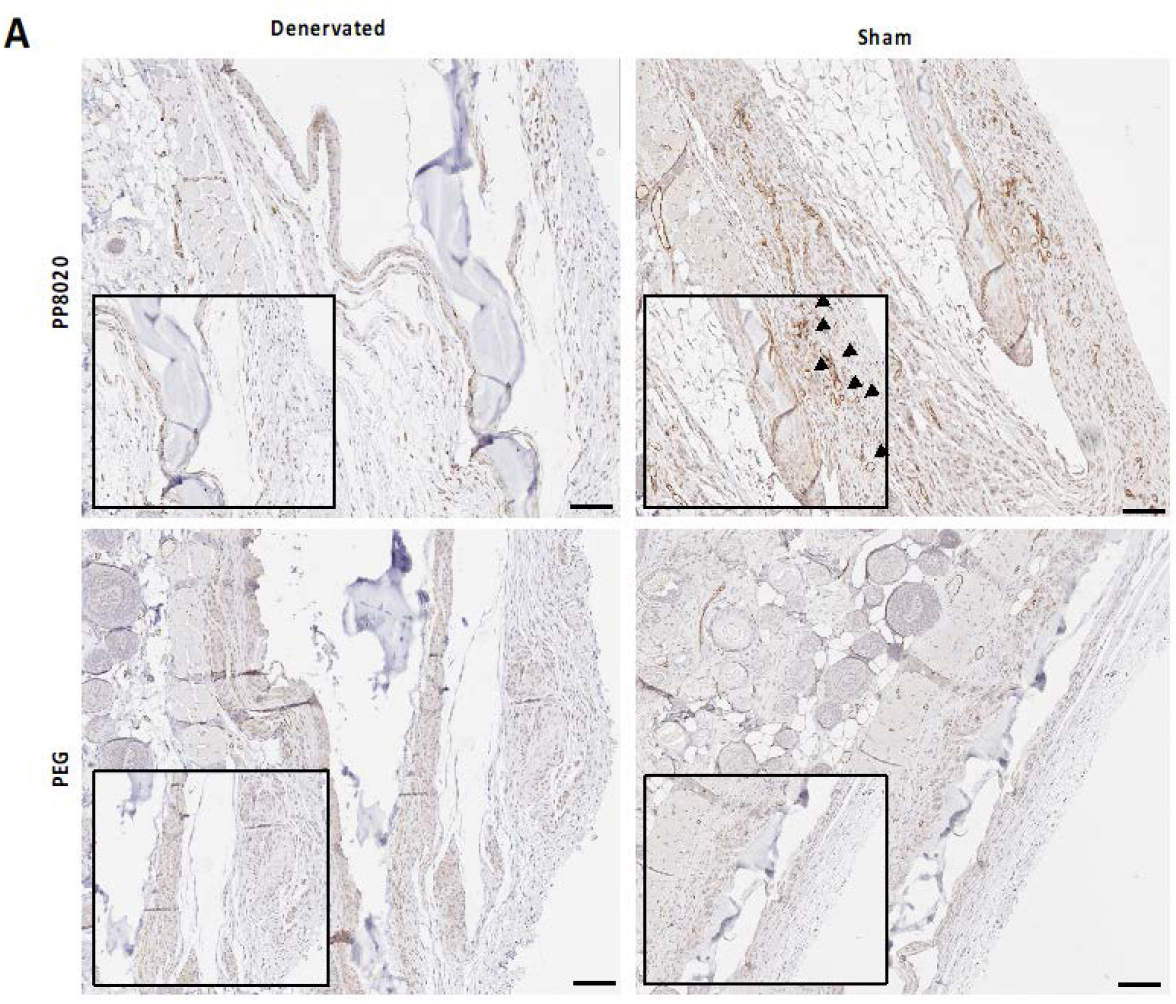
Innervation is required for methacrylic acid-induced angiogenesis. Denervation in subcutaneous tissue resulted in a decrease in the number of CD31 positive regions in 20% MAA explants relative to innervated sham operated control. PEG subcutaneous explants had fewer vessels. Representative CD31 stained histology sections 21 days after subcutaneous injection of PP8020 (20% MAA) or PEG hydrogels (arrows denote CD31 positive regions). Scale bar=100µm.

## DISCUSSION

### MAA-based biomaterials promote peripheral nerve growth in both CD1 and db/db mice

Our results highlight a new potential application of MAA-based biomaterials beyond exploitation of it’s vascular-regenerative properties. MAA-based biomaterials have been previously shown to promote angiogenesis[8,9] leading to its use in soft tissue repair[14,19] and islet transplantation[15,33]. Here we show that MAA-based hydrogels or a coating promote peripheral nerve growth. Implantation of MAA-coated silicone disks and hydrogels into the subcutaneous space of CD1 resulted in an increase in expression of peripheral nerve markers Uchl1 and βIII-tubulin at 21, 28, and 35 days relative to control materials. This increase in expression of nerve markers also coincided with an increase in nerves (PGP9.5 positive regions) in tissue treated with MAA, confirming that the increase in expression of nerve markers in tissues treated with MAA was due to nerve growth/increased nerve presence. Treatment of DRGs with MAA-coated biomaterials for 24 hours confirmed that MAA promoted greater axon outgrowth *ex vivo* compared to control materials. iDISCO-based three-dimensional imaging of MAA treated tissues further confirmed this, while intraplantar injections of MAA hydrogel exhibited a lower von Frey withdrawal threshold, indicating increased sensitivity, relative to PEG and contralateral controls. This increased sensitivity suggests that there are likely an increased presence of nerves in the MAA-treated paw. Importantly it appears that the nerve growth promoted by MAA is mechanosensitive, indicating that MAA generates nerve arborization/growth that is fully functional.

MAA-based biomaterials also increased the expression of neuronal markers and growth factors in a peripheral neuropathy model (db/db mice) over time relative to controls. This sustained increased in expression suggests that MAA may have the potential to prevent nerve damage in the context of injury wherein MAA prevents axon loss by upregulating key factors (such as Igf-1 and Shh) that promote axon survival. The development of a biomaterial that exploits MAA to prevent degeneration to complement a “regenerative” biomaterial may yield a promising new therapeutic avenue.

### MAA-induced peripheral nerve growth requires Igf-1 and Shh

Inhibition of *Igf-1* and *Shh* prevented MAA-induced peripheral nerve growth. While increased *Igf-1* has been shown to be macrophage-derived in an MAA-induced angiogenesis context[11] macrophage inhibition studies (clodronate liposomes) did not inhibit MAA-induced peripheral nerve growth, as demonstrated by the lack in the change in expression of nerve markers (*βIII-tubulin, Uchl1*, and *Shh*) compared to vehicle treated (PBS liposomes) mice (Supplementary Figure 2) following a 21 day inhibition study. This suggests that macrophage-derived growth factors might not have a central role in MAA-induced peripheral nerve growth. While macrophages have been shown to play a key role in MAA-driven vascularization at time-points as early as 7 days, MAA-induced axon growth occurs at 21 days. It’s likely that the increased levels of *Igf-1* and *Shh* observed following MAA implantation are derived from another cell type within the subcutaneous space, most likely peripheral nerve support cells such as Schwann cells or fibroblasts. Schwann cells possess the ability to dedifferentiate to a precursor state and secrete growth factors to promote nerve regeneration[2,34]. Moreover, ephrin-B/EphB2 signaling between Schwann cells and fibroblasts results in cell sorting and directional collective cell migration of Schwann cells to guide growing axons[35].

Denervation of tissues removes a necessary source of pro-regenerative paracrine signals which prevents/hinders tissue regeneration/repair, such as in the case of diabetic neuropathy. Here, denervation of the subcutaneous space prior to MAA delivery abolished the nerve growth effect. On the other hand, implantation of the MAA hydrogel into a diabetic neuropathy model was able to promote an increase in the expression of peripheral nerve markers. Unlike the complete denervation model, MAA was able to induce peripheral nerve growth in the diabetic neuropathy model. This is consistent with the idea that support cells (removed by denervation) may be key contributors to the beneficial impact of MAA. which is further evidenced in the denervation model in which absence of nerves hinders MAA-induced blood vessel generation.

The development of peripheral nerves relies on complex temporal and spatial signalling among axons, Schwann cells, and vessels[36]. Schwann cell proliferation is stimulated by neuronally-derived VEGF, which also promotes axonal outgrowth and the formation of microvessels[36]. Following complete nerve transection, Schwann cells utilize macrophage-derived blood vessels to guide regenerating axons back to the target tissue through a hypoxic VEGF-mediated response[37]. Physical or metabolic damage to peripheral nerves and Schwann cells results in transcriptional changes[38]. In the case of diabetes, hyperglycemia leads to an increase in oxidative stress, inflammation, and polyol pathway flux resulting in the impaired production of the Schwann cell-derived growth factors Shh, Ngf, and Cntf[39-41] ultimately leading to axonal degeneration and microvascular dysfunction[42]. Exogenous Shh and Igf-1 have been shown to reverse/mitigate the axonopathy associated with diabetes[27,40]. Use of the MAA hydrogel in diabetic db/db model resulted in an increase in the expression of Shh and Igf-1 as well as an increase in the expression of the peripheral nerve marker Uchl1 demonstrating that MAA is able to increase local trophic support for peripheral nerves as a mean to prevent further axonopathy and possibly promote regeneration.

### Functional Impact and Potential Applications

Innervation is essential for complete tissue repair. Peripheral nerves and supporting cells such as Schwann cells have been shown to mediate the regenerative capacity of tissues through the release of paracrine mediators from axon terminals[1-5] and through the regulation of tissue resident stem cell populations including hematopoietic and epidermal stem cells[43,44]. MAA-based biomaterials possess remarkable pro-regenerative capabilities, and here we identify a new neuro-regenerative application for these biomaterials. While the obvious application of MAA-based biomaterials in the context of peripheral nerve growth is its use in nerve conduits for large diameter nerve injuries, the more exciting application is the use of MAA to address the challenge of small diameter peripheral neuropathies not treatable through current nerve repair strategies, conduits and autografts[45]. Such neuropathies arise from trauma, autoimmune disorders, aging, metabolic disorders, or drug toxicity resulting in loss of motor, sensory, and autonomic functions and/or the development of weakness, numbness, and pain[46-48]. Current clinical treatment strategies aim to reduce pain, where a 30% reduction in pain is considered as successful[46-48]. No clinical treatment exists that target the underlying cause of peripheral neuropathies and promote nerve regeneration. Even with the gold standard of autografts for large diameter nerve injuries, full function of the nerve is rare[45]. Here we demonstrate that MAA-based biomaterials promote peripheral nerve growth in both a normal and neuropathic model and that the nerve growth induced by MAA is functional and capable of sensing and evoking a response to mechanical stimuli. Success of peripheral nerve regeneration is a complex multi-cellular process[49] dependent on a number of factors including the up-regulation of regeneration associated genes and growth factors, as well the establishment of an anti-inflammatory microenvironment permissive for growth. Several of these can be favourably manipulated/regulated by MAA, without the need for exogenous factors. Moreover, MAA is injectable, effective in small volumes, and is available as a degradable hydrogel for which the degradation rate can be tuned, making it a robust and versatile tool in the treatment of peripheral neuropathies.

Innervation is essential for all tissue engineering and regeneration strategies. The enhanced endogenous response driven by MAA-based biomaterials makes it an invaluable tool in enhancing the regenerative capacity of tissues of not only nerves but perhaps all complex tissues.

## ACKNOWLEDGMENTS

The authors acknowledge financial support from the University of Toronto’s Medicine by Design initiative, which receives funding from the Canada First Research Excellence Fund. A.M.A. was funded by an Ontario Graduate Scholarship. We thank Dr. Freda D. Miller and Matt Carr for their valuable advice and Dr. Lindsey Fiddes for technical assistance with microscopy. We would also like to thank Virginie Coindre for assistance with animal experiments.

## AUTHOR CONTRIBUTIONS

Alaura M. Androschuk: Conceptualization; Data curation; Formal analysis; Investigation; Methodology; Writing – original draft; Writing – review & editing

Michael V. Sefton: Conceptualization; Funding acquisition; Project administration; Writing – review & editing

Theresa H. Tam: Data curation; Investigation; Formal analysis

Redouan Mahou: Methodology

Chuen Lo: Investigation

Michael W. Salter: Project administration; Conceptualization; Methodology; Resources

## DATA AND MATERIALS AVAILABILITY

All data needed to evaluate the conclusions in the paper are present in the paper and/or the Supplementary Materials. Raw data available on request from corresponding author.

## SUPPLEMENTARY FIGURES

**Supplementary Figure 1.**
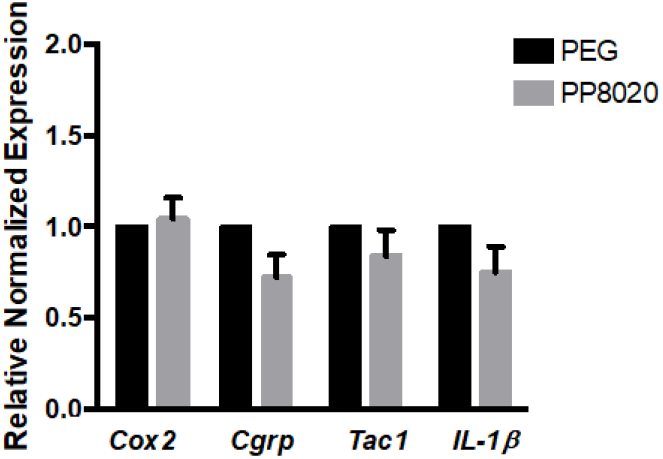
Methacrylic acid-based hydrogels did not increase expression of pain markers. 20% MAA injection did not increase the expression of the pain markers *Cox2, Cgrp, Tac1*, and *IL-1β* relative to PEG at day 21. Quantification of mRNA was performed as in Figure 1. and normalized to PEG samples. N=3 biological replicates per biomaterial group; error bars indicate S.E.M.

**Supplementary Figure 2.**
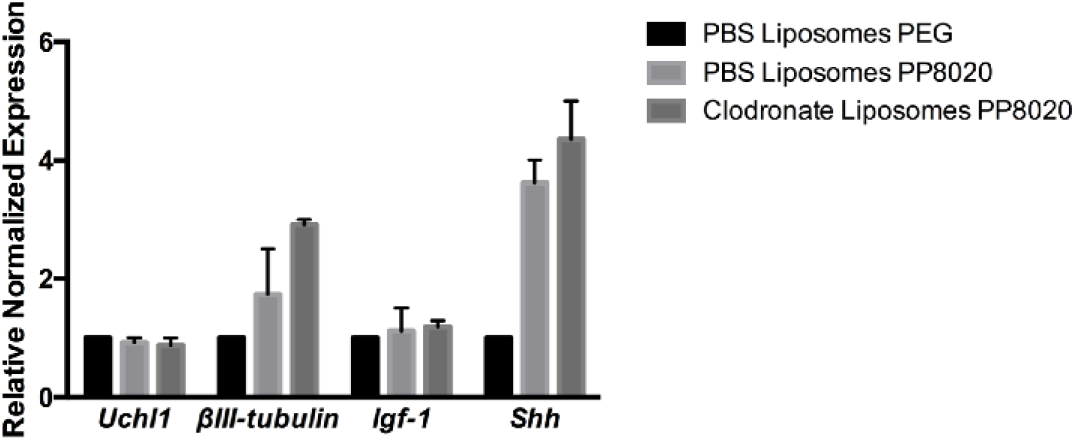
Macrophages are not directly responsible for MAA-induced peripheral nerve growth. Inhibiting macrophage recruitment with clodronate liposomes did not decrease the expression of *Uchl1, Igf-1*, or *Shh* in 20% MAA explants relative to PBS liposomes. qRT-PCR analysis of CD1 mice injected with PP8020 (20% MAA) and treated with the clodronate or PBS liposomes for 21 days. Quantification of mRNA was performed as in Figure 1. and normalized to PEG samples. N=2 biological replicates per biomaterial and treatment group; error bars indicate S.E.M.

